# The structural permissiveness of triosephosphate isomerase (TpiA) of *Escherichia coli*

**DOI:** 10.1101/2024.10.18.618993

**Authors:** Belen Calles, Borja Pitarch, Víctor de Lorenzo

**Author notes:** Correspondence to: Victor de Lorenzo National Center of Biotechnology CSIC Calle Darwin 3, Madrid 28049 (Spain) Pho.

## Abstract

Triosephosphate isomerase (TpiA) is widely regarded as an example of an optimally evolved enzyme due to its essential role in biological systems, its structural conservation, and its near-perfect kinetic parameters. In this study, we investigated the structural robustness of the archetypal TpiA variant from *Escherichia coli* using an in vitro 5- amino acid linker scanning method. The resulting library was introduced into a *tpiA* mutant strain for functional complementation. From this library, 15 TpiA variants that were phenotypically indistinguishable from the wild-type enzyme were selected for further analysis. Although all variants retained enzymatic activities within the wild-type range, several insertions were found in highly structured protein domains where the linker was expected to cause significant structural perturbations. Despite these potentially disruptive additions, the enzymes maintained their activity even when expressed in a *dnaK* mutant, suggesting that chaperones did not compensate for structural abnormalities *in vivo.* Additionally, when these mutant TpiA variants were produced using the PURE *in vitro* transcription/translation system, they exhibited enzymatic activity comparable to, and in some cases exceeding, that of the non-mutated enzyme. AlphaFold2 revealed that insertions reconstructed the local architecture of the nearby amino acid sequences. The evolutionary implications of this remarkable structural resilience are discussed.

## Introduction

Triosephosphate isomerase (TpiA) is a key enzyme of the standard glycolytic pathway that allows interconversion of the two products of fructose-1,6-biphosphate cleavage by Fda (i.e. dihydroxyacetone phosphate and glyceraldehyde-3-P) and thus the channeling of the whole C flow from glucose towards central metabolism ^[1]^. TpiA is one of the most conserved enzymes of the biological realm, not only in terms of activity and specificity, but also from a structural point of view ^[2]^. Inspection of homologous proteins borne by species covering the entire evolutionary tree indicates that the overall tridimensional organization of TpiA is basically kept all the way from archaea to humans ^[3]^. This conserved 3D architecture is the prototype of what has been called the TIM barrel folding pattern, which is adopted by enzymes with a wide variety of functions. The observed sequence similarity, or dissimilarity for that matter, has led to a debate regarding TIM-barrel evolution. Whether TIM barrels have resulted from convergent or divergent evolution remains an open question ^[4]^. The evolutionary history of this family of enzymes is still not completely certain, while some studies have suggested that there are evidences for both divergent ^[5]^ and convergent evolution ^[6]^, divergent evolution from a common ancestor explains more of the available data than does convergent evolution to a stable folding ^[7]^. Thus, the general consensus is that TIM barrels are divergently evolved from some ancestral protein This structure is common in metabolic biochemistry and some of the most effective enzymes belong to this category ^[8]^. The extraordinary maintenance of the 3D configuration (Fig. 1) could be an indication of an optimal evolutionary outcome to the biochemical problem of interconverting *in vivo* dihydroxyacetone phosphate (DHPA) and glyceraldehyde-3-phosphate (GA3P). TpiA is indeed a good example of a kinetically perfect enzyme, characterized by a very fast second-order constant, with a turnover that is limited only by the diffusion rate of substrate and enzyme ^[9]^, a feature considered an indicator of how far the catalyst has evolved ^[10]^. But such evolutionary optimum might be caused *in vivo* not only by inherent structural properties of the protein at stake in isolation. Much of the essentiality of given proteins stem from their capacity to interact with other molecular partners in the very crowded milieu of the cell cytoplasm and also their ability to respond to small molecules (e.g. allosteric regulation). These biological traits go beyond inherent structural stability and impose constraints that must be met to maintain the functioning of the corresponding biochemical or structural network. Not surprisingly, TpiA is also one of the most connected components —functionally and physically— of the proteomes of the organisms where the issue has been examined. In the case of *E. coli,* although the enzyme has no known interacting effectors other than its substrates, TpiA coalesces, whether by physical association or by functional interaction, with not less than 20 protein partners (Fig. 2). The evolutionary story of its structure is thus expected to reflect both a distinct individual robustness and a capacity to act together with other molecular companions. But how much is the contribution of each driving force i.e. *intrinsic* protein architecture for delivering the core enzymatic activity and *extrinsic* pressure for raising and maintaining productive contacts with other proteins? Surprisingly, virtually all platforms for structural prediction of protein configurations and domain flexibility rely on innate properties of the protein sequence with little or no consideration of such *in vivo* scenarios that can affect the most effective structure in the wider biochemical context of the cells.

**Figure 1.**
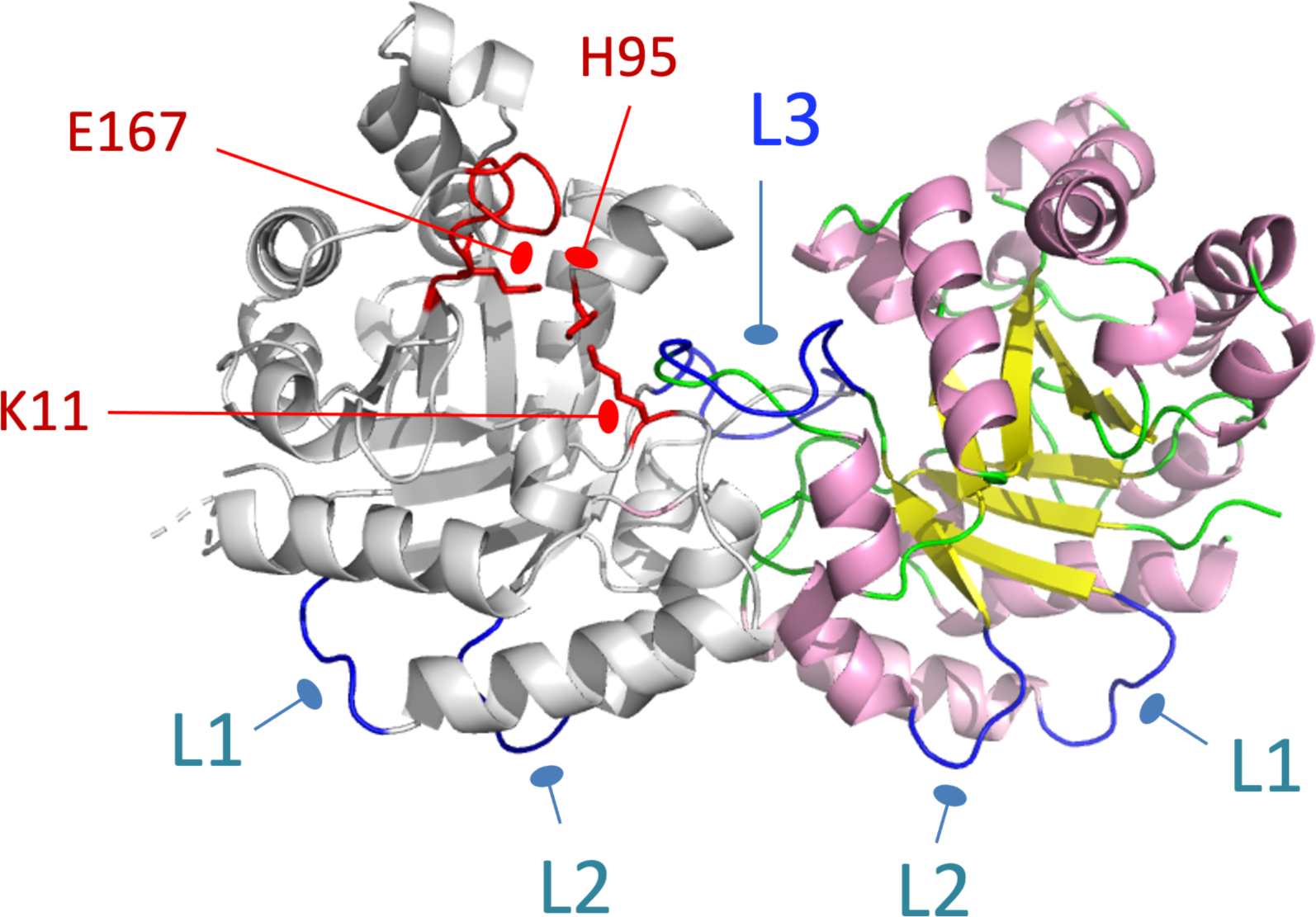
Structure of TpiA. The 3D architecture of *E. coli’*s triosephosphate isomerase (PDBj entries 1TRE and 4K6A) consists of two monomers, each formed by an eight-stranded αβ barrel, with one flexible loop region of ten amino acids containing the active site-lid at the C-terminal end of the barrel sheets (residues 165-175), including the key E167 residue that together with K11 and H95 are the three crucial catalytic residues for catalytic activity (lateral chains of the three main catalytic residues are shown in red). A number of possibly permissive loops represented as strings in blue.

**Figure 2.**
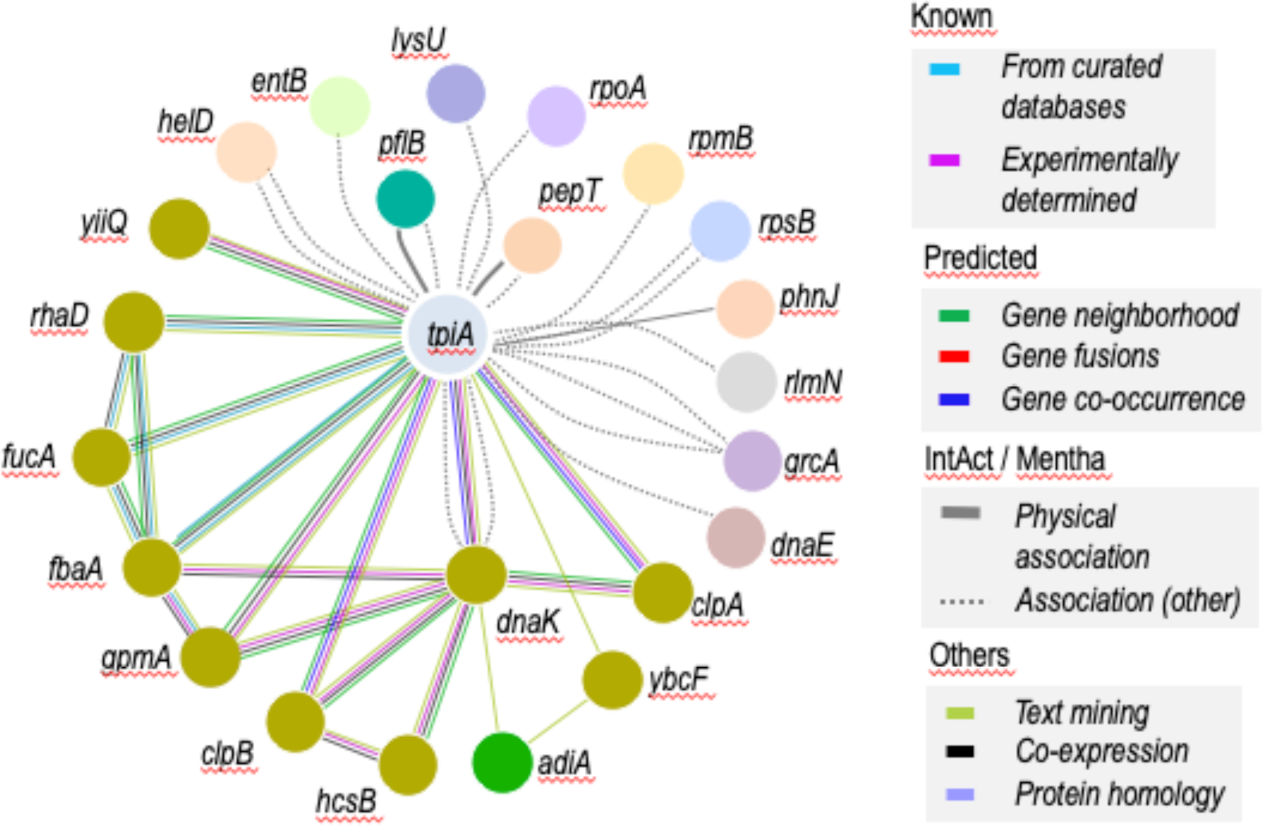
The TpiA interactome. The graph summarizes the interplay of TpiA with other cytoplasmic proteins retriever from DIP ^[39]^, Mentha ^[40]^, IntAct ^[41]^ and STRING ^[42]^). The figure shows a group of proteins known and/or predicted to interact according to different experimental and computational criteria ^[25, 43] [44]^. Interactions include both direct (physical as well as indirect (functional) associations—this does not necessarily mean they are physically binding to each other.

Most predictions are consistent with the notion that InDels (insertion-deletion) events are accepted by natural selection primarily if they occur in loops and turns, which themselves frequently occur on protein surfaces ^[11]^. From an evolutive perspective, it has been described that InDels accumulate in regions, which are generally more permissive to mutations, and specifically to loop regions much more frequently than in secondary structure elements ^[12]^. The idea that most, if not all, permissive sites locate in loop surface regions has been widely confirmed by experimental observations with specific proteins published to date ^[13]^, with the only exception of MalE protein, that probed to have a couple of permissive sites within structured regions, which can accept insertions of different lengths without loss of activity ^[14]^.

In this work we have taken advantage of the metabolic centrality of TpiA and the wealth of information about this enzyme for exploring the interplay between enzymatic activity and *in vivo* structural flexibility. *In silico* predictions of protein permissiveness are typically based on sequence conservation, homology models and thermodynamic calculations. In contrast, the work below takes a naïve approach to the same question by scanning the TpiA sequence with structurally disrupting peptides and selecting those that still keep full functionality *in vivo*. Such strategy for probing proteins by inserting foreign DNA sequences within the corresponding gene provides invaluable information about structure-function relationship ^[15]^. Linker scanning methods introduce sequences of different lengths into the target gene of interest which can then be mapped on physical or predicted tridimensional (3D) structures, thereby exposing the role of specific domains or motifs in either maintaining enzymatic activity or an interactive shape or both ^[16]^. As shown below, we probed the archetypal *tpi*A gene of *E. coli* for permissive locations in the encoded enzyme by using a Tn*7*-based transposon-mediated insertion scanning protocol that saturates every possible site of the targeted protein with 5-amino acid peptide segments that perturb secondary structures. The striking discrepancies between the permissive sites for such insertions predicted *in silico* and the ones selected *in vivo* (some of which hit highly structured protein locations) exposed an unexpected permissiveness of the protein to structural perturbations. These findings shed some light on the relative weight of endogenous physical architecture vs proteomic context for shaping the protein form that executes central metabolic reactions *in vivo*.

## RESULTS AND DISCUSION

### Benchmarking *tpi*A activity in a dedicated test strain

*Escherichia coli tpi*A gene encodes the only triosephosphate isomerase activity in this bacterium, which is a central enzyme of the carbon metabolism. *tpi*A is a stand-alone gene, likely to be expressed as a single cistron and transcribed independently of the adjacent genes (https://biocyc.org/gene?orgid=ECOLI&id=EG11015). On this basis we generated a deletion of the whole gene without affecting neighboring DNA sequences as explained in the Experimental Section, thereby producing *E. coli* W3110 Δ*tpi*A strain. Analysis of the knockout strain verified that, as expected, the *tpiA* gene is essential for cellular growth in glycerol, but not in glucose (not shown). This is because standard downwards glycolysis of the 6C-sugar can still proceed without interconversion of DHAP to GA3P. However, the loss of the enzyme impedes gluconeogenic synthesis of hexoses from 3C-compounds. The second component of the test system consisted of an E-tagged version of the gene/enzyme (*tpiA*ET). To this end, *tpi*A was cloned in an IPTG-dependent expression plasmid fused to a carboxy-terminal (C-terminal) E-tag epitope (see Experimental Section) to facilitate immunodetection and quantification of its expression levels *in vivo.* We then verified that this *tpiA*ET construct (borne by plasmid pBCL3; Fig. 3 and Table 1) could restore the growth of *E. coli* W3110 Δ*tpi*A to wild-type levels in minimal medium with glycerol as sole carbon source (Supplementary Fig. S1), indicating that the recombinant E-tagged TpiA protein (TpiAET) was fully active (see below). To further confirm this, we directly measured TpiA activity of both wild type *E. coli* W3110 and the Δ*tpi*A strain complemented with pBCL3. To do so, soluble fractions of each culture grown to stationary phase were analyzed by SDS-PAGE and the total protein contents were quantified with Bradford assays. Equivalent amounts of total soluble protein were used to perform enzymatic assays as explained in the Experimental Section. The complemented cells showed triosephosphate isomerase activity significantly higher than the wild type, consistently with the high levels of over-expression of TpiA in the thereby engineered strain (Fig. 3C).

**Figure 3.**
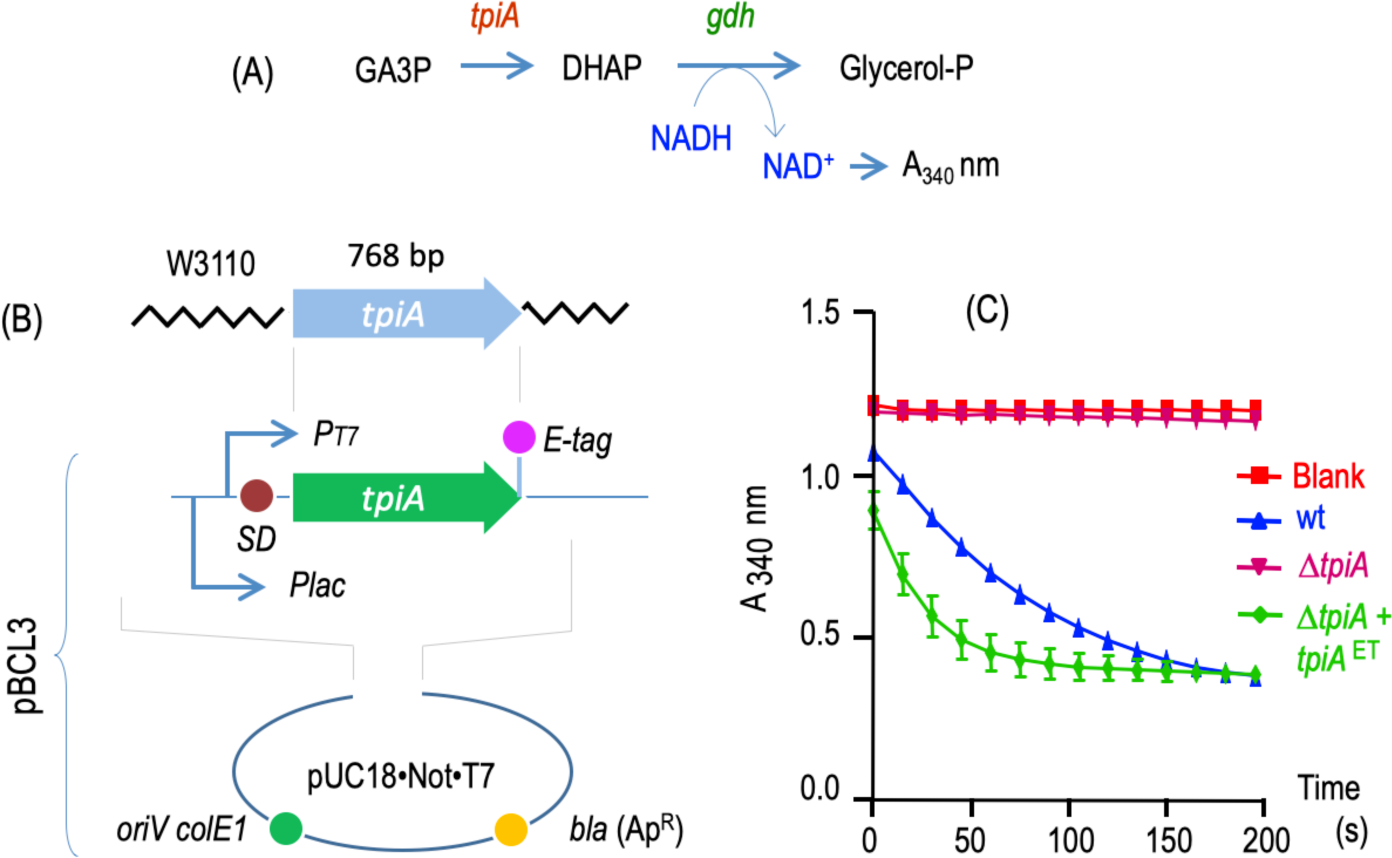
Components of the experimental system to probe structural resilience of TpiA. (A) Monitoring enzymatic activity of TpiA. As shown, NADH-dependent reduction of dihydroxyacetone phosphate (DHAP) resulting from TpiA catalysis to glycerol 3-phosphate can be spectrophotometrically measured at 340 nm, thereby reporting the previous step GA3P to DHAP. (B) Organization of expression plasmid pBCL3. The *tpiA* gene was amplified from the chromosome of *E. coli,* recloned in an expression vector (pUC18Not/T7; ^[30]^) preceded by a *Plac* promoter, a T7 promoter and a synthetic Shine-Dalgarno (SD) sequence for translation initiation. By default, *tpiA* gets expressed through the *Plac* promoter in pBCL3. (C) Reference enzymatic activity of wild-type TpiA and its E-tagged TpiA derivative (*tpiA*^EC^, green line). Note that the strain with the multicopy plasmid expressing *tpiA*^EC^ has more isomerase activity than the wild type, which carries just one chromosomal copy of the gene.

**Table 1.** Bacterial strains and plasmids used in this work.

| Strain/plasmid | Relevant features | Reference |
| --- | --- | --- |
| *E. coli* |  |  |
| W3110 | K12 F<sup>-</sup> IN (*rrnD-rrnE*) | Strain collection |
| W3110Δ*tpiA* | W3110 lacking the *tpiA* gene | this work |
| W3110Δ*dnaK* | W3110 lacking the *dnaK* gene | https://cgsc.biology.yale.edu/ |
| W3110Δ*dnaK-tpiA* | W3110 double *tpiA-dnaK* mutant strain | this work |
| XL1-Blue | *F'*[*proAB lacI<sup>q</sup> lacZ*ΔM15 *Tn10*] *endA1 hsdR17 supE44 thi-1 recA1 gyrA96 relA1, lac* | [37] |
| DHB4 | MC1000 *phoR* $\Delta$(*phoA*)*PvuII* $\Delta$(*malF*)3 *F'*[*lacI<sup>q</sup>ZYA pro*] | [38] |
| *Plasmids* |  |  |
| pUC18Not/T7<sup>*</sup> | pUC18Not with T7 promoter and *lacZp* in tandem; Ap<sup>r</sup> | [30] |
| pBCL1 | pUC18Not/T7<sup>*</sup> based vector inserted with a SD site between EcoRI and BanII sites | this work |
| pBCL2 | pBCL1 based vector added with an E-tag sequence inserted between PstI-HindIII sites | this work |
| pBCL3 | pBCL2 harboring *tpiA* gene cloned into SacI-BamHI sites | this work |

### Probing TpiA for structurally permissive sites

Perusal of the 3D organization of *E. coli’*s TpiA structures 1TRE and 4K6A deposited in PDBj databank (https://pdbj.org) and inspection of possible permissive sites based on sequence conservation using multiple alignment methods, pinpointed only 3 locations of the protein monomer predicted to have sufficiently flexible loops to accept peptide inserts without disrupting the rest of the protein edifice (Fig. 1). Loop 1 (L1) encompassed amino acids G32-G35 of the primary structure and loop 2 (L2) included the segment G56-H58. Finally, loop 3 (L3) spanned the chain V66-S71. The regions selected for permissive insertions should ideally show a low conservation in sequence, should not be involved in substrate binding, and should not be part of possible protein-protein interfaces. While L1 and L2 corresponded to poorly ordered, potentially malleable sequences in the tertiary structure of each of the monomers, L3 was too close to the monomer-monomer interface and was discarded as a possible target of implanting heterologous peptides. This was because changes in the L3 site could affect indirectly enzymatic activity even without disrupting organization of the monomer.

Using the homology modeling tool Modeller ^[17]^ for each loop region L1 and L2, a set of 100 loop models was generated based on the crystal structure. To evaluate the position most suited as insertion site, the Prosa’s energy function ^[18]^ was used. As a result, two specific insertion points, i.e. after A34 and S57 residues, were defined as optimal permissive sites in L1 and L2, respectively (J. Pleiss, personal communication). On these bases, laxness of sites A34 and S57 was tested experimentally by inserting genetically a heterologous 7 amino acid peptide (NVVVHQA) in each of the locations (thereby resulting in protein variants TpiaA34 and TpiAS57, respectively). The resulting carrier plasmids (pBCL3-A34 and pBCL3-S57; Table 1) expectedly complemented *E. coli* W3110 Δ*tpiA* for growth on glycerol (not shown). Once this was set, the next obvious question was whether structural flexibility TpiA was limited to these 2 recognizable sites or some additional plasticity could be endowed by the molecular milieu (other proteins, small molecules etc.) that may bind the protein *in vivo.* To uncover otherwise cryptic permissive sites of TpiA we used a transposon-mediated linker insertion scanning method (see Experimental Section). This is based on a mini-Tn*7* construct harboring an antibiotic resistance gene, flanked by *Pme*I sites at both ends ^[19]^. The *in vitro* process (sketched in Fig. 4A) involved transposition of the Tn*7* construct from the donor plasmid and its insertion into the target DNA cloned in a second plasmid, an action that happens essentially without sequence preference. The *in vitro* transposition reaction was then transformed in *E. coli* DHB4 electrocompetent cells and selected with Km (the antibiotic marker of the Tn7) and Ap, the antibiotic resistance of the target plasmid, to secure the recovery of inserted plasmids only. The whole plasmid content of the transformation pool was then extracted, digested with *Pme*I and religated for excising the bulk of the transposon. This leaves behind 10 base pairs that, together with the 5 pb duplication caused by transposon insertion, result in the occurrence of random 15 bp insertions through the entire length of the target DNA with the *tpiA* sequence. Two out of the three possible frames of the thereby implanted DNA are readable and thus bring about 5 amino acid-insertions when the target gene is translated—the third frame encodes a stop codon and therefore results in a truncated protein. Plasmid pBCL3 carrying the *tpiA*ET gene was subject to this procedure and then used as the target of the Tn*7* construct transprimer-5, which is delivered by pGPS5 vector (Table 1) through the action of an *in vitro* reaction driven by purified TnsABC* transposase. After selecting those insertions produced within the target gene a library of around 10^5^ *tpi*A derivatives was generated, each of which containing a 15 bp DNA insertion tagged with a unique *Pme*I restriction site—and two thirds of which should originate full-length enzyme variants.

**Figure 4.**
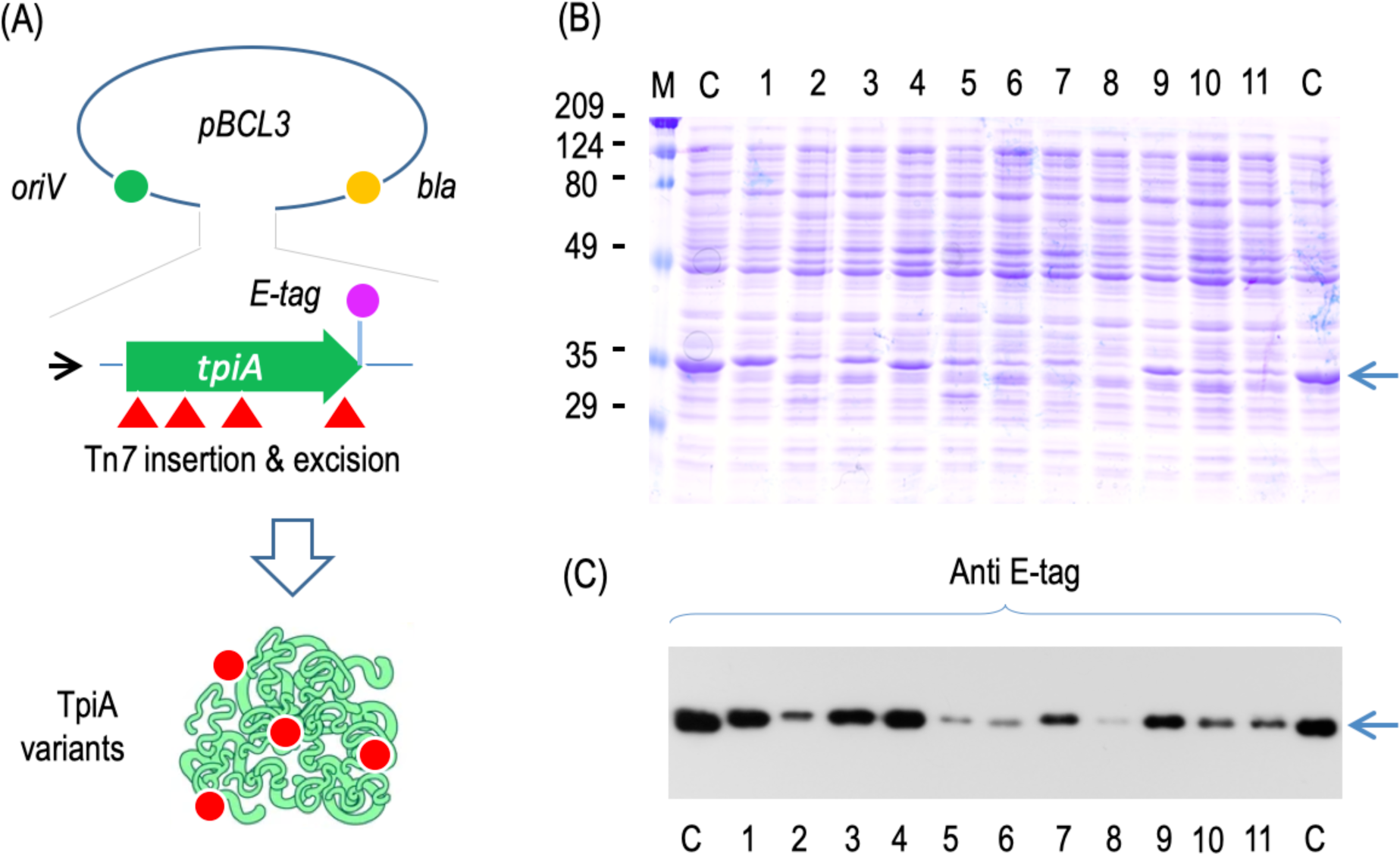
Generation of permissive pentapeptide insertions through the TpiA structure. (A) Strategy for generation of 5-aa inserts through the TpiA sequence. The transprimer (red arrow) delivered by the Tn7 transposition system was delivered to the *tpi*A gene cloned in pBCL3, in an *in vitro* reaction that was then transformed in a *tpi*A defective strain. Next, plasmids containing insertions within the gene of interest were digested with the unique PmeI site and religated, leading to a collection of *tpi*A gene variants, each of which harboring 15 bp DNA insertions that, one expressed, produced a comprehensive library of TpiA enzyme derivatives with inserts at different positions. (B) PAGE-SDS analysis of soluble protein fractions from clones bearing pentapeptide-inserted variants of TpiA and capable of complementing growth of a *E. coli* Δ*tpiA* mutant on glycerol. The arrow to the right indicates the band corresponding to TpiA. (C) Probing the gel above with an anti E-tag antibody with a Western blot assay. Note that the level of expression changes for each clone probably due to different protein solubilities.

### Analysis of peptide-scanned TpiA protein species

Following production of an insertion library in the *tpiA*ET we next pursued isolation of variants that still held a significant activity compared to the wild-type version. This was straightforward, as the loss of *tpi*A in *E. coli* W3110 makes bacteria unable to grow on minimal medium with glycerol as sole carbon source. Given that *tpiA*ET+ plasmid pBCL3 plasmid fully complements this mutant phenotype *in trans* (Supplementary Fig. S1) it became handy that the same complementation could be used for selecting permissive insertions through the TpiAET+ protein sequence. On this basis, the whole pentapeptide insertion library was transformed into *E. coli* W3110 *ΔtpiA* and plated on M9 glycerol medium with Ap. Approximately 10^3^ transformants were able to grow in such plates, with variable colony sizes from very small colonies to identical to the wild-type control (not shown). A number of healthy-looking clones were selected, separately grown in LB medium with ampicillin, the respective soluble fractions of the grown biomass of each culture obtained (see Experimental Section) and equivalent amount of protein loaded in SDS-PAGE and subject to immunoblotting with anti-E-tag antibodies. The same extracts were also subject to enzymatic assay to determine their triosephosphate isomerase activity. As shown in Fig. 5, while all clones complemented well the loss of *tpiA* in the host strain, there was not correlation between level of expression (compare with Fig. 4B and 4C) and level of activity, indicating that insertions had affected the protein in different ways. Just to illustrate this, variant #9 (Fig. 4) was expressed at the same level as the wild-type TpiAET protein but had a very poor activity (Fig. 5), while variants #2 and #5, which displayed a wild-type like enzymatic performance, were produced at lower levels than the wild-type protein in the soluble fraction. Only those variants showing an enzymatic activity ≥ 75% with respect to the intact TpiAET protein were selected as bearers of *bona fide* permissive insertions. For example, we picked TpiA-variants numbers #1 to #5, #7 and #11 but discarded #6 and #10, as activities were 50% or lower than the wild type (Fig. 5). In sum, 55 inserted and active clones were first checked but only 22 were considered entirely satisfactory TpiA variants. Table 2 lists the actual activities of these proteins in respect to the wild type enzyme with data normalized taking into account their differential expression as quantified by Western blot assays of Fig. 4C.

**Figure 5.**
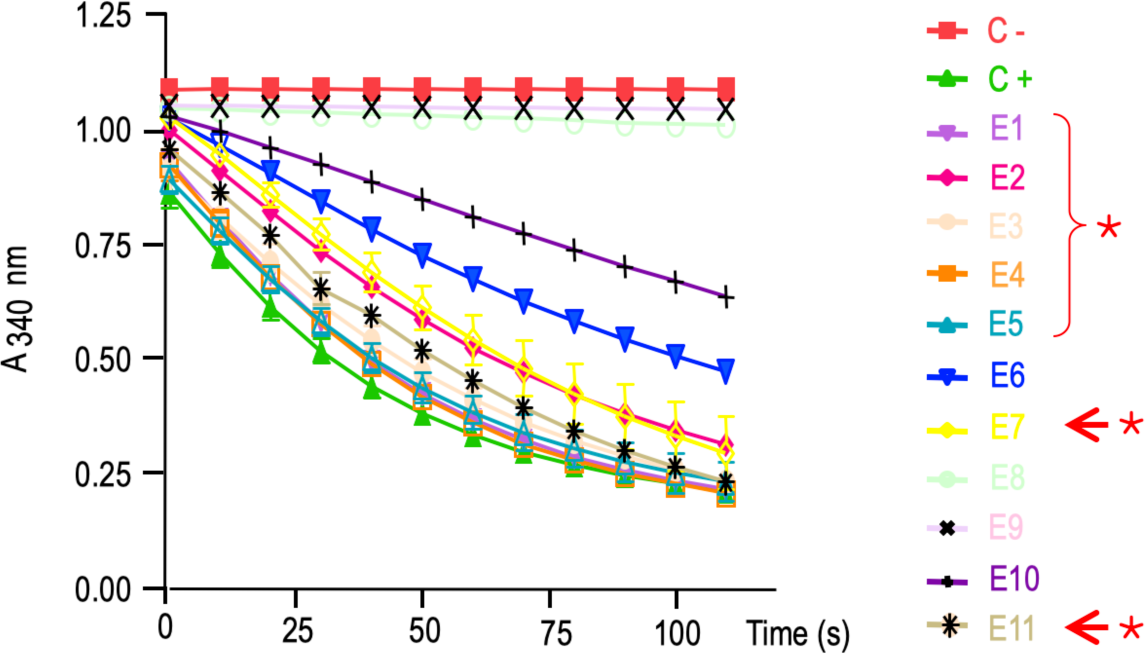
Triosephosphate isomerase activity of protein extracts from *E. coli* W3110 *tpiA* carrying pBCL3 derivatives encoding inserted variants of TpiA that complement the chromosomal mutation. Assays were run as indicated in the legends to Fig. 3C. Extracts from strains carrying either pBCL3 with the wild-type *tpiA* sequence or the insertless pUC18•Not•T7 vector were used as positive and negative controls C+ and C, respectively. The rest of the samples derive from the strains analyzed in Fig. 4B and 4C. Although all these clones complemented the growth phenotype in vivo, some of them showed a very reduced activity (e.g. E6, E8 and E9) and were not studied further. Others, however, had activities in the range of the wild-type (e.g. E1 to E5, E7, E11) and were picked for selected for sequencing and mapping of the permissive insertion sites (indicated with an *).

**Table 2.**
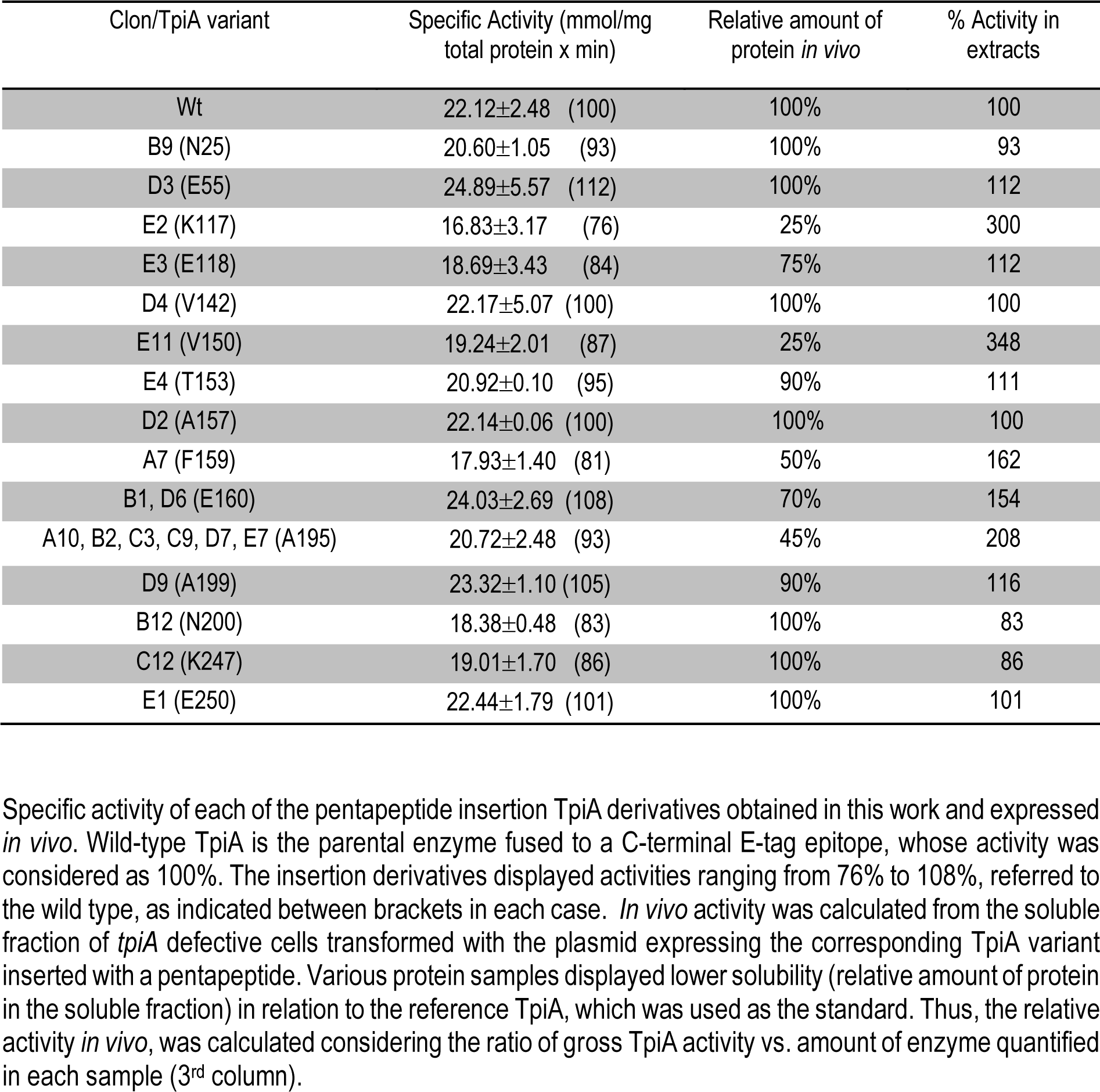
TpìA in protein extracts of E. coli W3110 *tpiA* expressing inserted variants of TpiA relative to their production levels *in vivo*.

### Localization of permissive insertions

All clones selected by their enzymatic activity were sequenced to locate the point of insertion. Sixteen different permissive sites were thereby mapped and found to be spread out along the entire length of TpiA protein (Fig. 6). Since we had selected only in-frame insertions, the result were five new amino acids within the TpiA protein that could be precisely localized in the crystallographic structure of the *E. coli* protein, which has been physically determined ^[20]^; PDBj entries 1TRE and 4K6A, respectively). Results of insertions into the known secondary and 3D structure of TpiA are summarized in Fig. 6 and 7. We found individual insertions after positions N25 in helix 1, E55 in loop3, K117 and E118 in helix 5; V142, V150 and T153 in helix 7; A157 and F159 in helix 8; A199 and N200 in helix 10; K247 and E250 in helix 13, A252 right after helix 13 in the C-terminal loop. In addition, various independent insertions were found both after residue E160 in loop 9 (two different clones) and also after A195 in helix 10 (where six independent insertions were localized). The majority of transpositions that spared the enzymatic activity of the protein, had occurred within highly structured regions of the TpiA architecture (see Fig. 6), including a few laying within α-helices. This was unexpected, as 5 amino acids insert more than one new turn (∼ 1.4 considering 3.6 aa per turn) within a pre-existing α-helix, yet they did not affect the functionality of the protein. It was also noteworthy that the predicted permissive insertion sites in the TpiA structure did not match the locations found with our random approach i.e. none of the permissive inserts mapped in L1, L2 or L3 Instead, various experimentally detected permissive spots lie inside in apparently structured enzyme regions (instead of loops) whose disruption is projected to kill protein function. Also of interest was that no insertions were found in any Δ strand. In the globular shape of the protein, Δ sheets form an eight-stranded Δ barrel surrounded by the alpha helixes, thereby suggesting that maintenance of Δ strands was essential for the activity of the enzyme. Finally, permissive insertions did occur only in polar protein spots of the protein dimer exposed to the external medium, excepting V142 which is partially hidden by the surrounding residues. This makes sense, as the hydrophobic core of any globular protein can hardly be dislocated without loss of function ^[21]^. But the question remained of how could activity be kept in full after the offsetting incorporation of 5 extra amino acids within well-structured parts of the enzyme.

**Figure 6.**
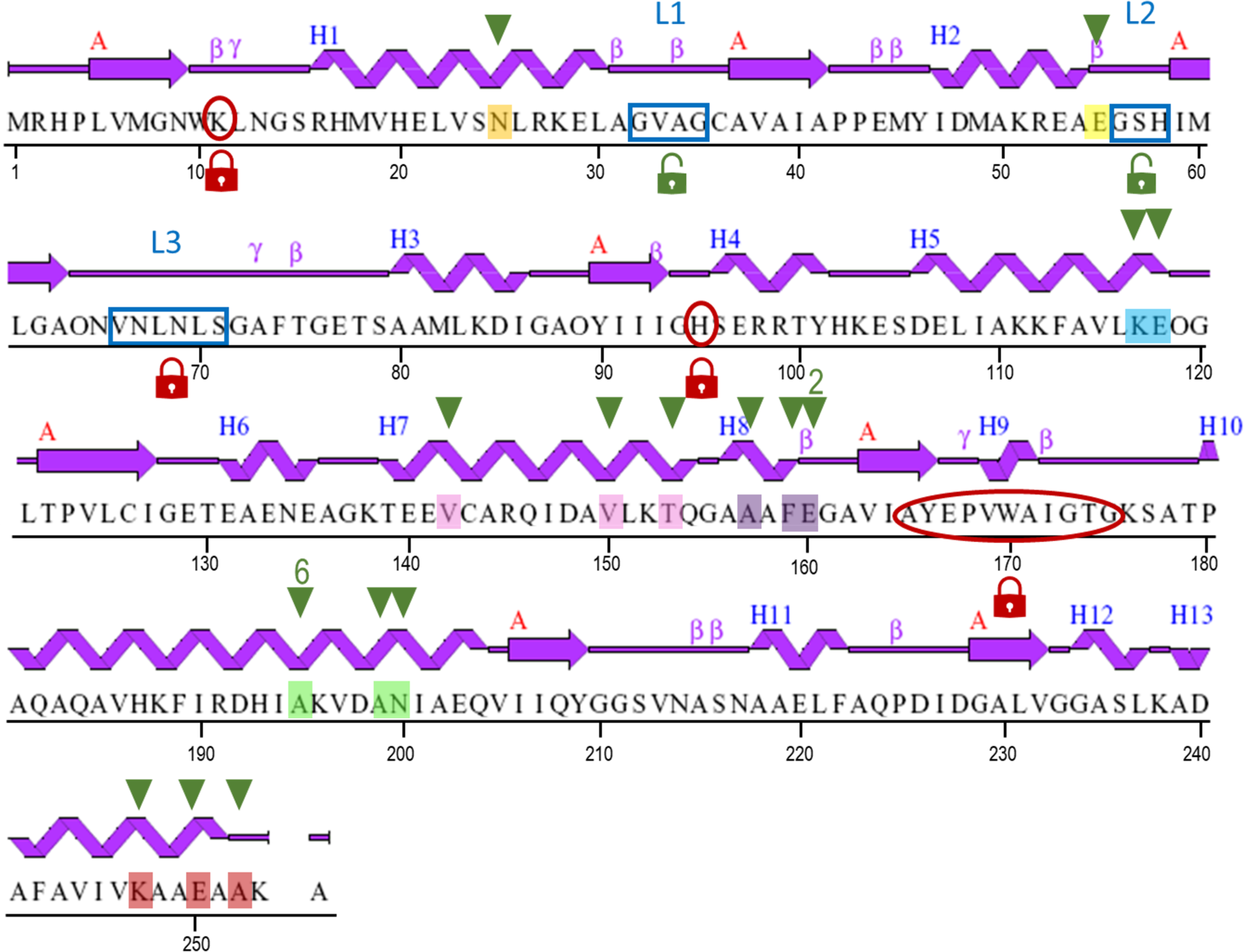
Distribution of permissive pentapeptide insertions trough the primary and secondary structure of TpiA. The secondary structure of the TpiA along its amino acid sequence was generated from 1TRE entry of PDBj data bank (https://pdbj.org/mine/structural_details/1tre), with an indication of α-helix and Δ-sheet motifs. Amino acids preceding permissive insertions are indicated with a color code that corresponds to the positions marked in the 3D structure of the enzyme in its dimeric active form shown in Fig. 7. Note that various independent insertions are located after E160 and A195 (2 and 6 events, respectively). Key residues of the active center of the enzyme are caged in open red circles, while unstructured loop regions L1, L2 and L3 are caged in blue. Red locks indicate essential residues that are not expected to be permissive while green locks mark putative permissive regions of the protein. Note that only one experimentally determined permissive site is located right before the L1 (after E55 residue, marked in yellow).

**Figure 7.**
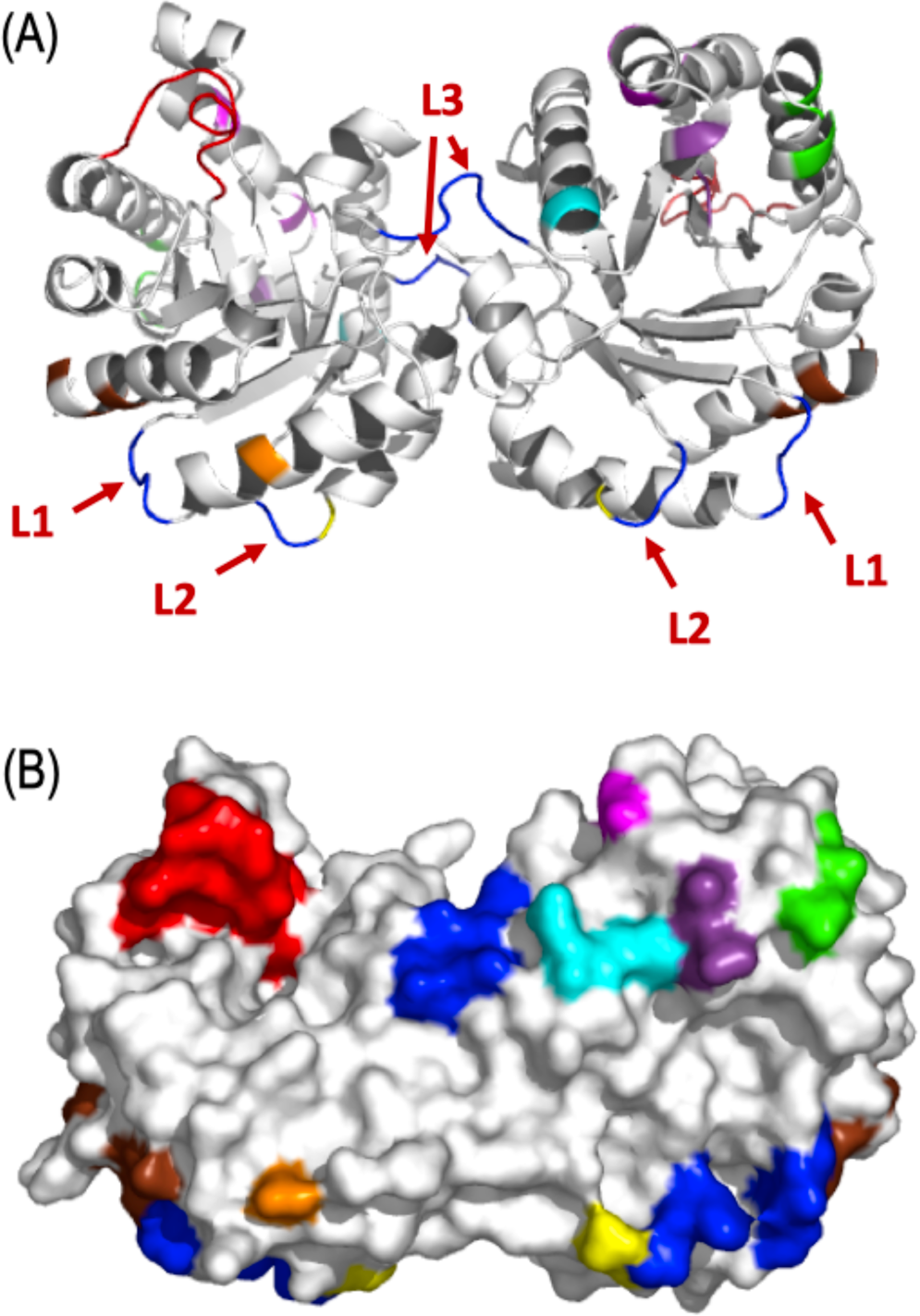
Spatial location of protein sites of TpiA hit by permissive pentapeptide insertions. (A) 3D String and ribbon structure of the TpiA dimer with an indication of potential (loops L1. L2, L3) and actual permissive sites. The location of the sites where permissive pentapeptide insertions were found are indicated with the same color code than those marked in Fig. 6. Amino acids that shape the active center are colored in red. Loops L1 and L2 (in blue) appeared upfront the best candidates for permissive insertions. Despite the lack of structure, L3 was not considered a location that tolerate inserts, as it lays at the dimerization interface. (B) Space-filling surface view of the TpiA homodimer highlighting relevant regions of the enzyme. The sites colored have the same code used in Fig. 6 and 7A: Loop regions (dark blue), active center (red) and permissive insertion sites (rest of colors). Most, if not all such spots were located in solvent-exposed locations.

### A role of DnaK chaperone in structural permissiveness of TpiA?

One avenue for explaining the tolerance of TpiA to insertions in surface-located α-helixes involves the stabilizing role of other interacting proteins *in vivo*. In particular, it has been shown before that chaperons of the Hsp70 family compensate structural disruptions caused by random mutations ^[22]^ thereby buffering the fitness cost of protein modifications ^[23]^, and facilitating protein evolution ^[24]^. DnaK is the bacterial homologue of Hsp70 and similarly to it, it is involved in the mechanism of formation of the tertiary structure of a nascent polypeptide chain preventing aggregation and promoting structuring to the native estate and even solubilizing and refolding of otherwise unstructured proteins. We thus hypothesized that the disturbance of TpiA by the inserted pentapeptides could be overcome through its combination with DnaK, which is indeed one of the proteins predicted to be associated with the enzyme (Fig. 2). To look into this possibility, we passed a representative subset of the plasmids bearing the TpiA insertion mutants to a *tpiA dnaK* host strain (Table 1) and inspected the growth of the individual transformants in minimal medium with glycerol as sole carbon source as a proxy of TpiA activity. The results are summarized in Table 3. Note that the growth of the double mutant with the plasmid encoding the wild-type *tpiA*ET gene without insertions loses approximately one third of its growth rate as compared to the isogenic *dnaK+* strain. This does not come as a surprise, as DnaK is involved in so many functions and its loss causes pleiotropic effects. When different TpiA variants were expressed in the *dnaK*- minus strain, growth rates in glycerol further dropped by 20-80 % with respect to the counterparts with the intact *tpiA*ET sequence. The highest impact of lacking *dnaK* was observed in insertions at residues K117, E119, V142 and V150, which happen to be the variants that yield less soluble protein fraction in the wild-type host strain (Fig. 4). While these results supported that DnaK helped maintaining an active structure of some of the mutated proteins, others bearing predictably disruptive insertions (e.g. A157, F159 and E160) remained unresponsive to the absence of the chaperon. The ensuing question was therefore whether this an in-built property of the proteins at stake or other interacting *partners in vivo* (possible carried on in the derived extracts) could be responsible for the maintenance of an active structure.

**Table 3.** Effect of *dnaK* on *in vivo* activity of pentapeptide-inserted TpiA variants.

| TpiA Variant | *tpiA* Strain |  | *tpiA-dnaK* Strain |  |
| --- | --- | --- | --- | --- |
| C + (wt) | 0.199 ± 0.003 | (100 %) | 0.071 ± 0.074 | (100 %) |
| C - ($\Delta tpiA$) | 0.013 ± 0.002 | (6 %) | 0.012 ± 0.004 | (17 %) |
| N25 | 0.168 ± 0.017 | (85 %) | 0.049 ± 0.001 | (68 %) |
| E55 | 0.200 ± 0.025 | (101 %) | 0.077 ± 0.003 | (108 %) |
| K117 | 0.163 ± 0.012 | (82 %) | 0.035 ± 0.002 | (49 %) |
| E118 | 0.151 ± 0.029 | (75 %) | 0.012 ± 0.006 | (18 %) |
| V142 | 0.159 ± 0.024 | (80 %) | 0.030 ± 0.003 | (42 %) |
| V150 | 0.119 ± 0.017 | (60 %) | 0.015 ± 0.004 | (30 %) |
| T153 | 0.176 ± 0.010 | (89 %) | 0.054 ± 0.007 | (76 %) |
| A157 | 0.154 ± 0.010 | (78 %) | 0.073 ± 0.004 | (102 %) |
| F159 | 0.143 ± 0.018 | (72 %) | 0.079 ± 0.001 | (110 %) |
| E160 | 0.134 ± 0.016 | (68 %) | 0.064 ± 0.004 | (91 %) |
| A195 | 0.141 ± 0.018 | (71 %) | 0.036 ± 0.005 | (51 %) |
| A199 | 0.150 ± 0.003 | (76 %) | 0.044 ± 0.002 | (61 %) |
| N200 | 0.140 ± 0.026 | (70 %) | 0.040 ± 0.004 | (57 %) |
| K247 | 0.171 ± 0.003 | (86 %) | 0.052 ± 0.005 | (73 %) |
| E250 | 0.161 ± 0.010 | (81 %) | 0.044 ± 0.003 | (61 %) |

Growth rates of TpiA insertion derivatives in the presence and in the absence of DnaK chaperone. Cultures of *tpiA*-defective and *tpiA dnaK* double null strains were transformed with plasmids expressing the corresponding TpiA variants, to monitor their growth capacity. Plasmids with the wild-type *tpiA* version as well as the empty vector were included as positive and negative control, respectively. The growth rate of cultures harboring the wild type TpiA were considered as 100% in both genetic backgrounds. The growth rates of the rest of TpiA derivatives were normalized by the respective wild-type data for easy comparison (% between brackets in both mutant strains).

### *In vitro* production and properties of TpiA variants

As shown in Fig. 2, triosephosphate isomerase possesses a quite populated interactome which, apart of DnaK, includes central carbon metabolism enzymes, nucleic acids and a number of other, diverse functionalities. Considering that proteins often function as part of a complex network, including direct physical interactions ^[25]^, we wondered whether the structural tolerance of TpiA stems from its own constitution or depends on the assistance of other molecular partners. To look into this, we synthesized *in vitro* selected TpiA variants with insertions spanning the entire protein structure using a minimal reconstituted transcription-translation platform (PURExpress® system, see Experimental Section). Next, we examined the activities of the thereby generated proteins and compared them to those measured in soluble cell free extracts (see Fig. 9). The results showed a general decrease in the activity of the TpiA variants with respect to that obtained from the *in vivo* experiments. The activity drop ranged from 20 to 80 per cent, suggesting that factors absent in the *in vitro* reaction, could be facilitating TpiA activity. These differences suggested a role of host factors (missing in the *in vitro* system) in the production, folding and activity of some of the inserted protein variants (e.g. N25, K117 and V142), but also exposed that other mutants (e.g. E55, E160 and A195) inherently kept a virtually full catalytic capacity in the absence of any other aid which could assist folding and/or enzymatic performance.

**Figure 8.**
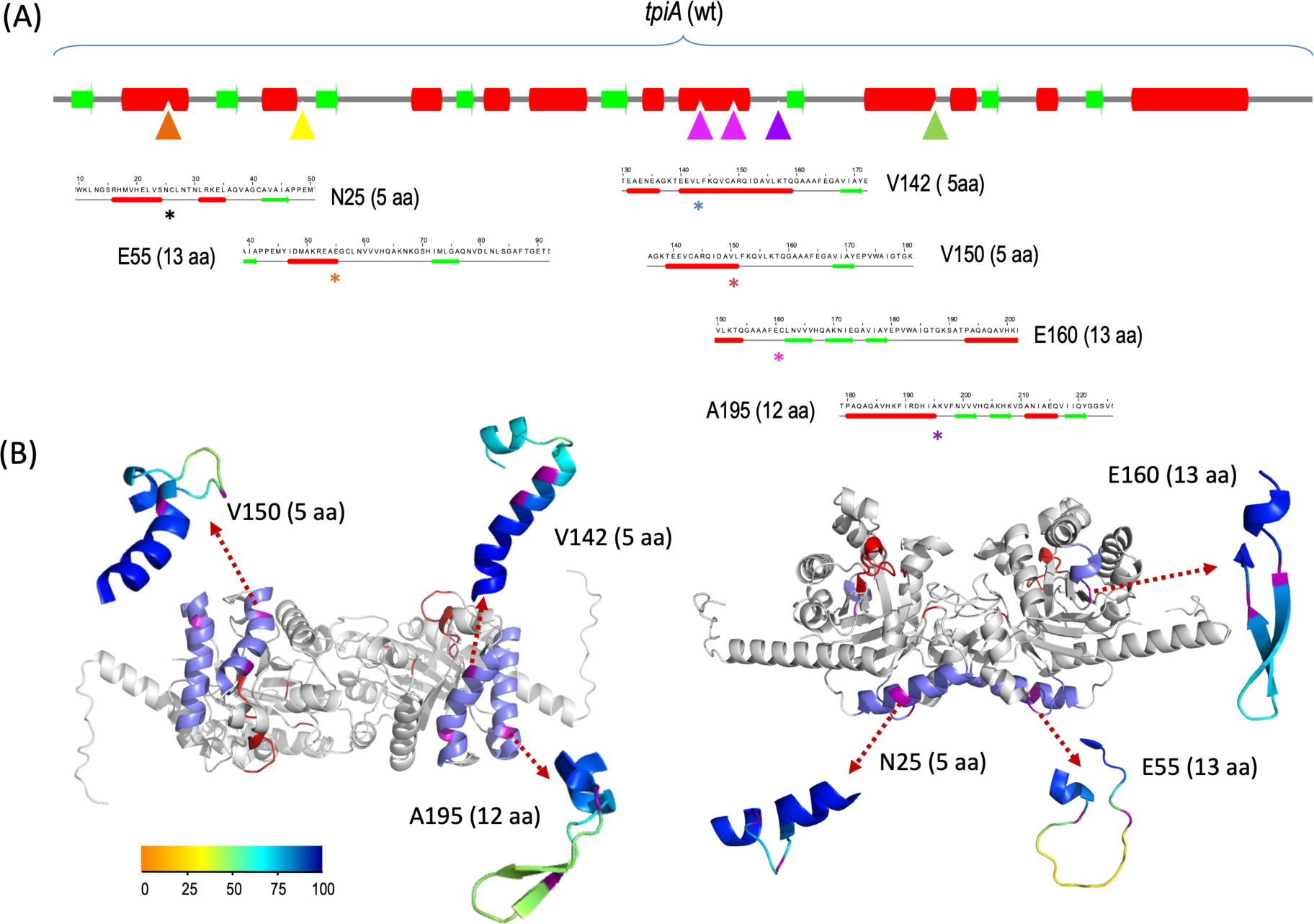
Structural consequences of pentapeptide insertions in permissive locations of the 3D TpiA architecture. (A) Secondary structures of TpiA protein and 6 different inserted derivatives predicted by JPred (https://www.compbio.dundee.ac.uk/jpred/). Helixes are marked as red rods, and sheets as green arrows. The sites of individual insertions in the wild-type sequence are marked with triangles with the same color code as the asterisks in the regions affected in each of the insertion derivatives shown beneath. Only changes in the near vicinity of the point of insertion of the corresponding segment are shown, as none of them affected the structure of the rest of the protein. (B) 3D structure of the TpiA enzyme and its insertion derivatives, modelled by AlphaFold2 (https://alphafold.ebi.ac.uk/) and represented in cartoon format. The structure of the wild type protein fused to a C-terminal E-tag, is shown in light grey color, except those regions affected by different insertions, which are blue colored and marked with the exact point of insertion in magenta. Key residues for the catalytic pocket have been colored in red. For each of the independent insertions shown in this figure, changes in the protein structure have been represented in the blowups, which show the putative folding of the region. In every blowup, amino acids marked in magenta stand for the starting and end point of the corresponding inserted peptide, while the rest of amino acids added up are colored on a PLDDT basis as shown in the corresponding color bar scale. Note that the length of peptide insertions ranges from 5 to 13 amino acids, depending on the specific insertion, as denoted in each case.

**Figure 9.**
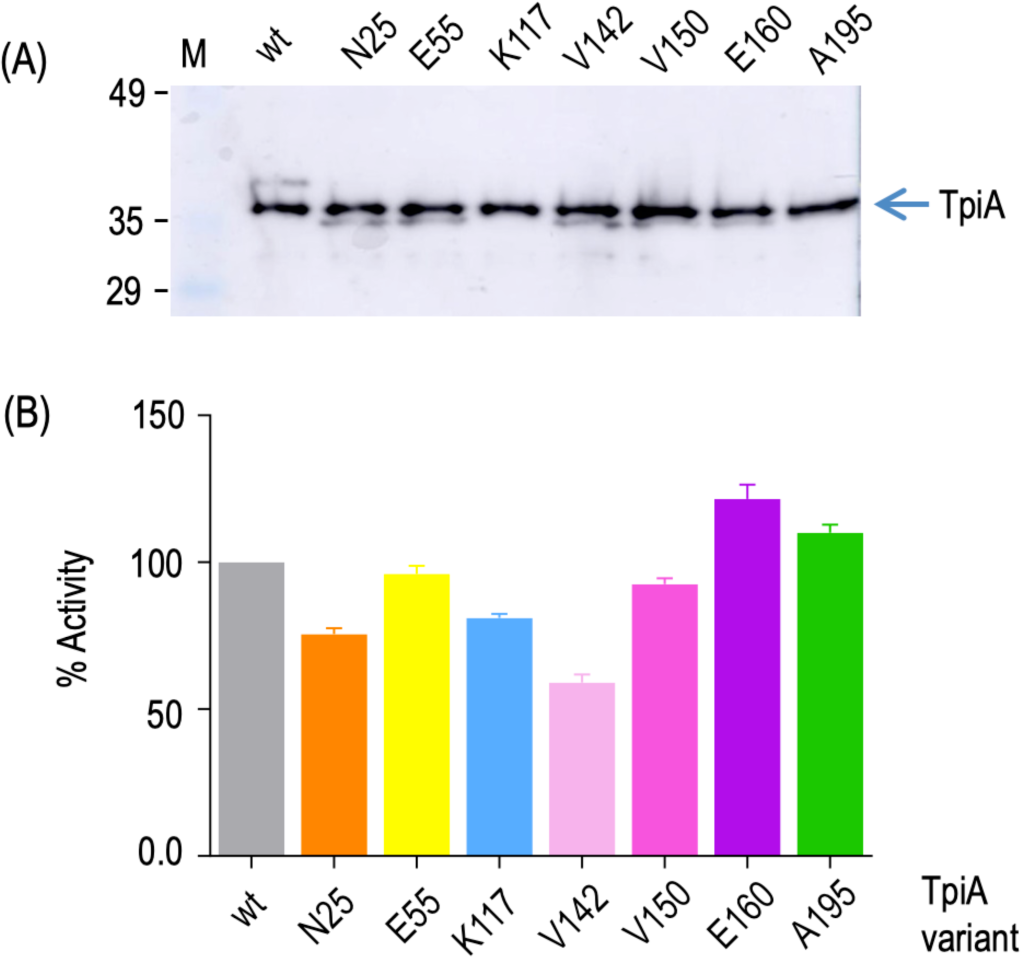
Expression and activity in vitro of TpiA variants bearing pentapeptide insertions in permissive sites. (A) *In vitro* expression of selected TpiA derivatives in a PURExpress® protein synthesis platform (see Experimental Section). Samples run in a PAGE-SDS gel TpiA^ET^ variants detected in in a Western blot assay with anti-Etag antibodies. The corresponding insertion points within the secondary structure of TpiA enzyme are indicated on top. (B) Enzymatic activities of TpiA variants synthetized *in vitro.* The reference wild type activity was then considered 100 per cent and the others normalized to this value.

### Probing permissiveness to longer amino acid insertions

On the basis of the above we set out to explore whether some of the protein sites more tolerant to inserts of the pentapeptide generated with the Tn7-based method could also accommodate progressively longer implants. To this end, we picked 3 different permissive positions to replace the 5-amino acid insets by a stretch of seven residues (NVVVHQA), which were added by site directed mutagenesis as explained in the Experimental Section. Location E55 was the only permissive site found experimentally that locates close to one of the predicted permissive loops (L2). Spots E160 and A195 were picked because both of them were found more than once in the initial screening. Specifically, insertion at E160 were found twice and lies at the beginning of a short loop. Finally, position A195, which was found six times during the opening survey, is located in the middle of the 25 aa α-helix-rich P182-Q206 of the protein structure. The new insertion variants were subject to functional complementation tests *in vivo* and enzymatic activity assays as before. As shown in Supplementary Fig. S2, extending the length of the inserts from 5 to 7 amino acids had no impact on TpiA performance *in vivo* and *in vitro*, as the corresponding parameters were indistinguishable from those of the wild type TpiAET protein. To further survey the ability of these regions of the protein to accept even longer amino acid additions, we took advantage of the *Pme*I site located located at the 15 bp insertion left by the transposition procedure to add DNA sequences either coding for additional 8 residues (total insertion of 39 bp, i.e. 13 aa long, in the case of E55 and E160 locations) or 7 residues in A195 position (total insertion size 36 bp, i.e. 12 aa long). Every new construct with the extended insertion was able to complement the TpiA null strain and gave enzymatic values similar to that of the wild type protein (Supplementary Fig. S2), indicating that the regions tested are highly permissive to rather long insertions without significant loss of activity.

### AlphaFold2 Modelling

To understand the rationale of how TpiA can accommodate insertion peptides within the different permissive sites described in this work, including relative long peptides in highly structured positions, without losing its enzymatic properties, we modelled the structure of all the inserted TpiA variants described above, by using the Alpha-Fold2 platform, which produces highly accurate predictions ^[26]^. To this end, the wild type TpiA protein, containing the C-terminal Etag sequence, was first aligned with one of the crystal structures available at the PDB (4K6A), confirming that they completely overlapped (excepting the extended C-terminal region where the E-tag epitope was added, not shown). Analysis of each of the variants constructed and described in this work, including site directed insertions into the putative optimal permissive loops, pentapeptide random insertions, along with longer insertions described in the preceding section, showed that the overall protein shape as well as the integrity of the active center of TpiA was not affected in any case. Most pentapeptide inserts did not disrupt conserved structural motifs but instead reconstructed the local architecture of the nearby amino acid sequences (see, for example, insertions V142 and V150 in Fig. 8). Only at position N25, within the alpha-helix 1, the pentapeptide insertion produced a small unstructured loop that precedes the rest of the alpha-helix, that is predicted to refold after such perturbation (Fig. 8A and 8B).

The structure of a set of longer insertions introduced into specific permissive sites of the protein, that did not decrease the enzymatic activity of TpiA as reported above, were also predicted by AlphaFold2. As observed with smaller five-residue insertions, the general 3D structure of the enzyme was neither affected in this case. However, inserts in position E160 and A195, located in structured regions of the protein, are predicted to form beta-hairpin motifs, while the one after E55 remains as a random coil region (see Fig. 8). Eight of the amino acids added (NVVVHQAK) to lengthen the pentapeptide insert at these three sites are shared by the three constructs under consideration (see the specific sequence of each of the insertions in Supplementary Table S1). The fact that Δ strands hairpin motifs are only predicted by AlphaFold2 when the insert locates into a structured region of the protein, i.e. at positions E160 and A195, but not when inserted into a loop, i.e. after E55 residue, suggests that there is a strong influence of the local surrounding structural motifs that determines the outcome of the insert. Yet it seems clear that sequence determinants of β-hairpin structure stability also make a significant major contribution, as described elsewhere ^[27]^.

## Conclusion

The work presented in this article documents the structural resilience of the glycolytic enzyme triosephosphate isomerase (TpiA) from *E. coli* by demonstrating that TpiA is highly tolerant to a range of predictably disruptive perturbations. Despite the insertion of five-amino acid linkers into various parts of the enzyme, including very structured regions, the protein retained significant activity *in vivo* and *in vitro.* This indicated that the protein is remarkably robust and can accommodate alterations without losing functionality. A distinct feature of TpiA exposed by the data above is that permissiveness of some protein variants seems to be an inherent property of the protein sequence, rather than being dependent on external factors such as molecular chaperones and accompanying interactome—as some TpiA variants maintained activity even when produced entirely by themselves. This may reflect the evolutionary history of the enzyme, as its ability to maintain function despite structural changes suggests a high degree of optimization, including tolerance to perturbations, which may contribute to its widespread maintenance across different species. Finally, our results also raise a word of warning that predictions based solely on *in silico* models might not fully capture the structural flexibility observed *in vivo.* In this respect, this study emphasizes the importance of experimental validation, as the observed elasticity of TpiA was not predicted by computational models. In sum, we argue that these findings provide insights into the balance between intrinsic structural properties and external factors in maintaining enzyme function, with potential implications for understanding protein evolution and stability.

## Experimental Section

### Bacterial strains, media and general methods

The strains and plasmids used in this work are listed in Table 1. The parental wild-type strain used was strain W3110 of *E. coli* K-12. The derivative *tpiA* knock-out strain was constructed by introducing a kanamycin (Km) resistance gene in place of the chromosomal *tpiA* gene ^[28]^. After selection of a clone inserted with the Km gene -flanked by FRT sequences, the antibiotic resistance was eliminated by FLP-mediated excision with pCP20 helper plasmid, which was afterwards cured by growth at 37°C. The *tpiA dnaK* double mutant strain was also constructed with the same method, using the *dnaK* strain (obtained from the *E. coli* Genetic Stock Centre) as template. Briefly, the Km resistance gene inserted in *dnaK* strain, together with homology regions flanking the gene (143 bp upstream and 153 bp downstream *dnaK* gene), was amplified by PCR using oligonucleotides dnaK-F y dnaK-R (see Supplementary Table S2). The resulting DNA fragment was electroporated into competent cells of the *tpiA* mutant harboring the *red* recombinase expression plasmid pKD46. Km resistant colonies were checked by PCR using appropriate primers to detect the replacement of *dnaK* gene.

The selected clone was also re-checked for *tpiA* deletion. Then, the helper plasmid pKD46 was cured by growth at 37° C. *E. coli* XL-1 Blue and DHB4 strains were used for general cloning procedures. Solid and liquid growth media were either LB or M9 minimal medium supplemented with 15 µM thiamine and 1% glycerol as the sole carbon source. When required, ampicillin (150 µg/ml) or kanamycin (75 µg/ml) were added. Standard cloning procedures were used as described ^[29]^.

### Construction of plasmid pBCL3

The construct of reference for this work was built by inserting a DNA segment encompassing a canonical Shine Dalgarno site, an E-tag sequence and the coding region of the *tpiA* gene in vector pUC18Not/T7* ^[30]^. For this, we hybridized oligonucleotides rbs-F and rbs-R, flanked by EcoRI (5’) and BanII (3’) tails, including the RBS sequence as well as XhoI, BglII, EcoRV and StuI restriction sites. The resulting DNA fragment was ligated to pUC18Not/T7* digested with EcoRI and BanII producing plasmid pBCL1. Next, hybridization of oligonucelotides Etag-F and Etag-R gave rise to a DNA fragment coding for the E-tag sequence with PstI/HindIII tails, which was ligated into the same sites of pBCL1 resulting in pBCL2. In parallel, genomic DNA was extracted from *E. coli* W3110 and used as template (100 ng) to amplify the *tpiA* gene in a buffer containing 2 mM MgCl2, 0.5 µM of primers tpiA-F and tpiA-R, 125 µM of deoxynucleosides triphosphate mix (dNTPs) and 3 u of Pfu DNA polymerase (Promega). Reaction was run with a first step of 4 min of denaturalization at 94°C, followed by 30 cycles of denaturalization (94°C, 1 min), annealing (65°C, 1 min) and polymerization (75°C, 1 min) with a final step of extension (75°C, 10 min). The amplification fragment was then digested with SacI and BamHI restriction enzymes and ligated into the corresponding sites of pBCL2 finally producing the *tpiA*ET -expressing plasmid pBCL3.

### Generation of peptide-insertion mutants in *tpiA*

The target gene *tpiA,* cloned in pBCL3, was subject to an *in vitro t*ransposition experiment using the linker scanning mutagenesis system, according to the supplieŕs instructions (GPS^LM^-LS Linker-Scanning System, New England Biolabs). The insertion of the Tn7 based transprimer into the target plasmid produced two classes of hybrid plasmids containing the transprimer in either the vector backbone or in the target gene. To select only target DNA insertions, this pool of plasmids was digested with SacI and XbaI, generating four different restriction bands, which were separated in a 0.8% agarose gel. After isolating and purifying the band consisting of the target DNA with the transprimer insertion the DNA obtained was recloned in pBCL2 digested with the same enzymes. This cloning produced a library of Tn7 insertions in the *tpiA* gene. Digestion of the plasmid DNA prepared from this library with PmeI yielded two fragments: the bulk of Tn7 transprimer and linearized plasmids. Then, the purified band corresponding to linearized plasmids was religated to produce a library of cloned tpiA genes, each of which containing a 15 bp insertion. Expression of this library produced a collection of TpiA proteins harboring randomly located pentapeptide insertions. Site-directed 7-amino acid insertions in selected positions of *tpiA* gene were done following Quick-Change^TM^ method modified by Wang and Malcolm ^[31]^, using oligonucleotides A195F and A195R (for insertion after A195 residue), E55F and E55R (for insertion after E55 aa) and E160F and E160R (for insertion after position E160). Longer insertions were done by inserting a dsDNA assembly produced from hybridized oligonucleotides tailored for this purpose, into PmeI sites present into each original pentapeptide insertion. Note that *Pme*I restriction site cuts within one particular codon when insertion occurs in one frame but between two different codons in the second frame while third frame gives rise to a stop codon and thus produces truncated proteins, which were not selected in our analysis.

### Cell culture and protein extracts

To generate enough biomass for production of extracts, cells were grown at 37°C in LB medium supplemented with Ap, with vigorous shaking to an OD_600_ of 0.4-0.5 and expression of the triosephosphate isomerase was induced by addition of IPTG (100 µM) for three hours. Then, cells were harvested by centrifugation at 7,000 x g for 10 minutes at 4°C. Cells chilled on ice were disrupted by sonication in seven 15 s periods alternating with 1 min periods of cooling in ice. The crude cell lysate was centrifuged at 12,000 xg for 30 min at 4°C. Protein concentration of the supernatant was measured by Bradford assay ^[32]^.

### Protein electrophoresis and western blot analysis

Soluble cell extracts were separated by SDS-PAGE electrophoresis (10%) and later Coomassie blue-stained and/or electroblotted onto a PVDF Inmobilon-P membrane (Millipore). To specifically detect TpiAET, HRP-anti E-tag conjugate antibodies (Pharmacia) were used at a final dilution of 1:5000. Detection of soluble fractions was carried out with a luminiscence reaction using a solution of 1.25 mM Luminol (Sigma), 40 mM Luciferin (Roche) and 0.0075% (v/v) hydrogen peroxide (Sigma) in 100 mM Tris- ClH pH 8 or with BM Chemiluminiscence Blotting Substrate (POD, Roche). After 1 min incubation in the dark, blots were exposed to X-ray films (X-OMAT, Kodak). Where needed, signals in the membranes were directly captured and integrated using the software package included in an Amhersam Imager 600 System (General Electric).

### Enzyme assays

Triosephosphate isomerase activity was followed as the change of absorbance at 340 nm due to the oxidation of NADH in a coupled-enzyme assay based on the method of Plaut and Knowles ^[33]^. The conditions for the forward reaction forming dihydroxyacetone phosphate were 100 mM TEA pH 7.9, 5 mM EDTA pH 8, 0.14 mM NADH (Roche), 0.4 mM DL-glyceraldehyde-3-phosphate, 2 µg/ml of glycerol 3-phosphate dehydrogenase (Roche) and 5 µg of cell free protein extract, in a final volume of 1 ml. All enzyme assays were initiated by the addition of the substrate and were performed at room temperature (22 °C). The decrease of A_340nm_ was recorded for two minutes in a spectrophotometer Ultrospec 3000 pro (Amersham Pharmacia Biotech), except in the experiment shown in supplementary Fig. S1, were reactions (final volume of 200 μl) were performed in microtiter plates and recorded with a Victor-2 plate reader (Perkin Elmer) Every experiment was done with two biological samples and three technical replicates.

### *In vitro* protein synthesis

A number of plasmids encoding wild-type TpiA or its derived pentapeptide-inserted variants were selected from the opening library and separately used as template for PURExpress® In Vitro Protein Synthesis assays (New England BioLabs inc.) according to the supplier instructions. TpiA variants were transcribed from the T7 polymerase promoter located upstream of each of pBCL3 derivatives carrying the cognate sequences (Fig. 3B). Template preparation was done with phenol:chloroform extraction followed by ethanol precipitation to remove nucleases (DNases and RNases) that could be inhibitory for the transcription/translation process. Reactions were assembled on ice by mixing 10 µl of solution A, 7.5 µl of solution B, 10.3 units of ANTI-RNase (Ambion^TM^), 300 ng of template plasmid and nuclease-free H_2_O to a total volume of 25µl. Then, the mixtures were incubated at 37°C for two hours after which the reactions were stopped by placing the tubes on ice. Enzymatic TpiA activity of the reaction products was then analysed and normalized by the relative amount of TpiA protein quantified by western blotting analysis of the *in vitro* reaction products using anti E-tag antibodies and the Amersham Imaging AI600 system as before.

### Analysis of secondary and tertiary structure of TpiA insertion variants

All predicted structures were generated using AlphaFold2 ^[34]^ v2.2.0 multimer mode, including the wild type, as it contained a tag in the C-terminal region. We checked the secondary structure of the insertions using JPred Secondary Structure Prediction Web Service ^[35]^ integrated in Jalview tool, as well as the secondary structure proposed by Alphafold2, also visualized using Jalview ^[36]^. For visualization, we used Pymol https://pymol.org and, for the structures with insertion, we used a spectrum colour scale based on the pLDDT value similar to the one in the AlphaFold Database website https://alphafold.ebi.ac.uk/ as shown in Fig. 8.

## Data availability

The data that support the findings of this study are available from the corresponding Author upon reasonable request

## Acknowledgments

Research in VdL Laboratory is funded by the NYMPHE (HORIZON-CL6-2021-UE 101060625) Contract of the European Union, the BIOSINT-CM (Y2020/TCS- 6555) Project of the Comunidad de Madrid - European Structural and Investment Funds (FSE, FECER) and Grant CEX2023-001386-S of the MICIU/AEI. Authors thank Sven Panke (ETH Zürich) for inspiration and many suggestions in the earlier stages of this work. We are indebted to Jürgen Pleiss (Department of Chemistry, Universität Stuttgart, Germany) for data on the *in silico* search for permissive sites as well as for helpful discussion relating some of the *in vivo* permissive sites found in this work. We also thank Florencio Pazos (Systems Biology Department, National Center of Biotechnology CSIC, Madrid, Spain) for support through this project and critical reading of the manuscript.

## Notes

### Competing Interest Statement

The authors have declared no competing interest.

