## Supplementary for "The structural permissiveness of triosephosphate isomerase (TpiA) of *Escherichia coli*"

### Supplementary Information to Calles *et al.*

#### The structural permissiveness of triose phosphate isomerase (TpiA) of *Escherichia coli*

**Supplementary Figure S1.** Verification of triose phosphate isomerase activity *in vivo* of cloned and E-tagged *tpiA*

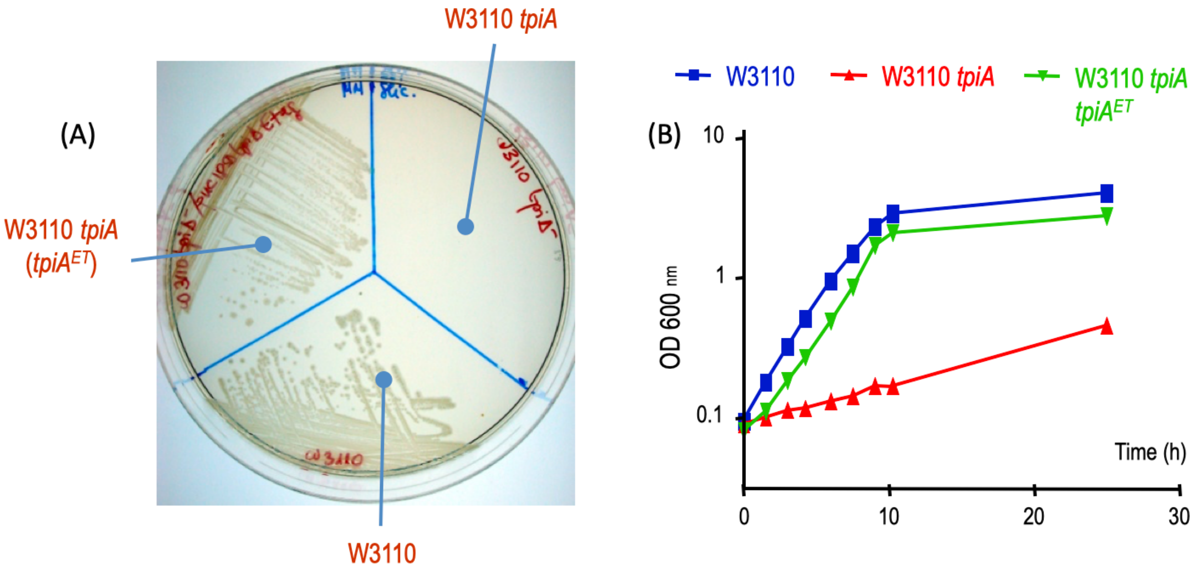


(A) Complementation assays. *E. coli* W3110 *tpiA* transformed with *tpiA*$^{ET+}$ plasmid pBCL3 was streaked on minimal medium with glycerol as sole carbon source along with positive (wild type W3110) and negative (plasmid-free W3110 *tpiA*) controls. (B) Growth curves of the same strains in M9 medium with glycerol as sole C source.

**Supplementary Figure S2.** Characterization of TpiA variants inserted with >5 amino acids at selected permissive sites.

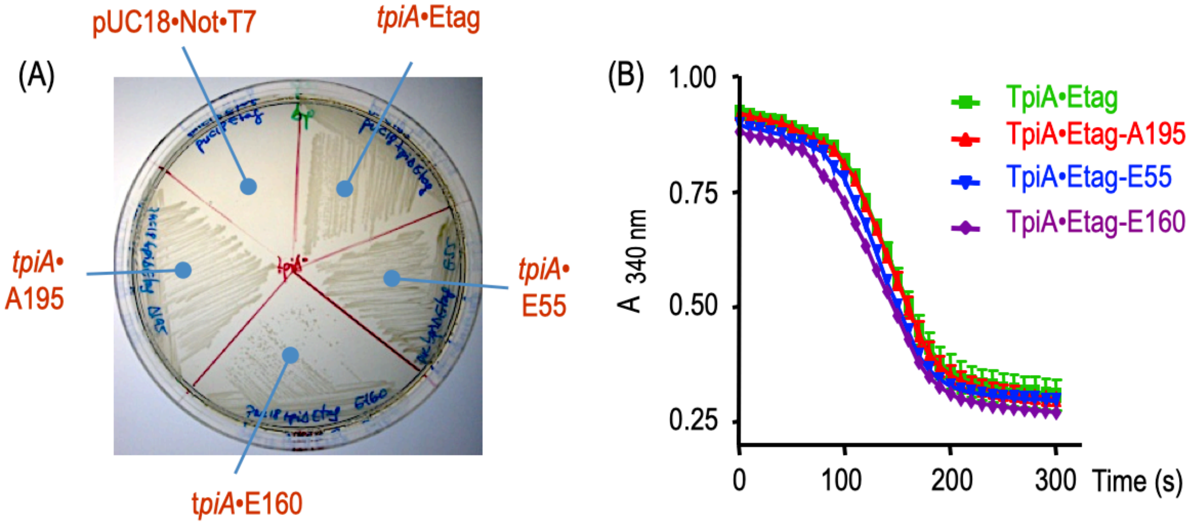


(A) Complementation assays. Plasmids expressing different TpiA species harboring peptides longer than five amino acids at three different positions of the protein were transformed in *E. coli* W3110 *tpiA* and plated on minimal medium with glycerol as sole carbon source as indicated along with positive (plasmid expressing wild-type *tpiA*) and negative (vector alone) controls. (B) Triose phosphate isomerase activity of extracts derived from the same strains.

**Supplementary Table S1.** Plasmids containing TpiA insertion variants constructed in this work.

| **Insertion plasmids constructed in this work** | **Relevant features (point of Insertion in *tpi*A gene)** | **Insertion Sequence** |
|---|---|---|
| pBCL3-N25 | Pentapeptide insertion after N25 residue | ...22-VSN**CLNTN**LRK-29.... |
| pBCL3-A34 | Heptapeptide insertion after aminoacid A34 (loop 1, L1) | ...31-GVA**NVVVHQA**LRK-38.... |
| pBCL3-S57 | Heptapeptide insertion after aminoacid S57 (loop 2, L2) | ...54-EGS**NVVVHQA**HIM-61.... |
| pBCL3-E55(1) | Pentapeptide insertion after E55 residue | …52-EAE**GCLNK**GSH-59 |
| pBCL3-E55(2) | Thirteen aa insertion after E55 residue | …52-EAE**GCLNVVVHQAKNK**GSH-59… |
| pBCL3-K117 | Pentapeptide insertion after K117 residue | …114-VLK**VFKQK**EQG-121… |
| pBCL3-E118 | Pentapeptide insertion after E118 residue | …115-LKE**LFKQE**QGL-122… |
| pBCL3-V142 | Pentapeptide insertion after V142 residue | …139-EEV**LFKQV**CAR-146… |
| pBCL3-V150 | Pentapeptide insertion after V150 residue | …147-DAV**LFKQV**LKT-154… |
| pBCL3-T153 | Pentapeptide insertion after T153 residue | …150-LKT**CLNKT**QGA-157… |
| pBCL3-A157 | Pentapeptide insertion after T153 residue | …154-GAA**CLNTA**AFE-161… |
| pBCL3-F159 | Pentapeptide insertion after F159 residue | …156-AAF**CLNTF**EGA-163… |
| pBCL3-E160(1) | Pentapeptide insertion after E160 residue | …157-AFE**GCLNK**GAV-164 |
| pBCL3-E160(2) | Thirteen aa insertion after E160 residue | …157-AFE**CLNVVVHQAKNIE**GAV-164… |
| pBCL3-A195(1) | Pentapeptide insertion after A195 residue | …192-HIA**KVFKH**KVD-199… |
| pBCL3-A195(2) | Twelve aa insertion after E160 residue | …192-HIA**KVFNVVVHQAKH**KVD-199… |
| pBCL3-A199 | Pentapeptide insertion after A199 residue | …196-VDA**CLNNA**NIA-203… |
| pBCL3-N200 | Pentapeptide insertion after N200 reside | …197-DAN**CLNTN**IAE-204… |
| pBCL3-K247 | Pentapeptide insertion after K247 reside | …244-IVK**VFKHK**AAE-251… |
| pBCL3-E250 | Pentapeptide insertion after E250 residue | …247-AAE**VFKQE**AAK-254 |

The specific insertion sequences are denoted in bold-blue

**Supplementary Table S2.** Oligonucleotides used in this work

| Oligonucleotide | Sequence |
|---|---|
| rbs-F | AATT*GAATTC*CTCGAGAGATCTGATATCAGGAGGCCTGAGCT |
| rbs-R | CAGGCCTCCTGATATCAGATCTCTCGAG*GAATTC* |
| Etag-F | GGCCGCAGGTGCGCCGGTGCCGTATCCGGACCCGCTGGAACCGCGTTAA |
| Etag-R | AGCTTTAACGCGGTTCCAGCGGGTCCGGATACGGCACCGGCGCACCTGCGGCCTGCA |
| tpiA-F | AACAC*GAGCTC*ATGCGACATCCTTTAGTGATGGG |
| tpiA-R | ACGC*GGATCC*AGCCTGTTTAGCCGCTTCTGCAG |
| dnaK-F | ACGTGGTTTACGACCCCATTTAG |
| dnaK-R | CTGATTTCACGCTCTTCCGCTG |
| A195F | CATCCGTGACCACATCGCTAACGTGGTGGTGCATCAGGCGAAAGTTGACGCTAACATCGC |
| A195R | GCGATGTTAGCGTCAACTTTCGCCTGATGCACCACCACGTTAGCGATGTGGTCACGGATG |
| E55F | GATATGGCGAAGCGCGAAGCTGAAAACGTGGTGGTGCATCAGGCGGGCAGCCACATCA TGCTGGG |
| E55R | CCCAGCATGATGTGGCTGCCCGCCTGATGCACCACCACGTTTTCAGCTTCGCGCTTCGC CATATC |
| E160F | CAGGGTGCTGCGGCATTCGAAAACGTGGTGGTGCATCAGGCGGGTGCGGTTATCGCTT ACGAAC |
| E160R | GTTCGTAAGCGATAACCGCACCCGCCTGATGCACCACCACGTTTTCGAATGCCGCAGCA CCCTG |

Restrictions sites used for cloning purposes are marked in underlined-cursive characters
